# REEP4 is recruited to the inner nuclear membrane by ELYS and promotes nuclear pore complex formation

**DOI:** 10.1101/2020.09.30.320333

**Authors:** Banafsheh Golchoubian, Andreas Brunner, Helena Bragulat-Teixidor, Busra A. Akarlar, Nurhan Ozlu, Anne-Lore Schlaitz

**Affiliations:** Center for Molecular Biology of Heidelberg University, Heidelberg, Germany; Department of Molecular Biology and Genetics, Koç University, Istanbul, Turkey

**Keywords:** nuclear envelope, endoplasmic reticulum, mitosis, reticulon homology domain

## Abstract

Nuclear pore complexes (NPCs) are channels within the nuclear envelope that mediate nucleocytoplasmic transport. NPCs assemble either into the closed nuclear envelope during interphase or concomitantly with nuclear envelope reformation during anaphase. Both, interphase and post-mitotic NPC biogenesis require local deformation of membrane. Yet, the factors that control proper membrane remodeling for post-mitotic NPC assembly are unknown. Here, we report that the reticulon homology domain-protein REEP4 localizes not only to high-curvature membrane of the cytoplasmic endoplasmic reticulum (ER) but also to the inner nuclear membrane (INM). We show that REEP4 is recruited to the INM by the NPC biogenesis factor ELYS and promotes NPC assembly. REEP4 contributes mainly to anaphase NPC assembly, suggesting that REEP4 has an unexpected role in coordinating nuclear envelope reformation with post-mitotic NPC biogenesis.

## Introduction

Nuclear pore complexes (NPCs) are essential gateways for nucleocytoplasmic transport and are built from multiple copies of around 30 different proteins called nucleoporins, or Nups (Hampoelz et al., 2019). Nucleoporins assemble at the nuclear pore, an opening created by fusion of the two lipid bilayers of the nuclear envelope. The walls of the nuclear pore form a highly curved domain within the otherwise flat nuclear membrane. NPCs and the nuclear envelope disassemble during prophase of mammalian open mitosis, releasing soluble Nups into the cytosol and membrane-bound proteins into the ER (Güttinger et al., 2009). During anaphase, ER-localized inner nuclear membrane proteins bind to the segregated chromosomes and initiate nuclear envelope reformation (Anderson et al., 2009). Concurrently, a step-wise process reconstructs NPCs (Schellhaus et al., 2016). The NPC biogenesis pathway that initiates during anaphase is referred to as post-mitotic NPC assembly and depends on the nucleoporin ELYS. ELYS binds to chromatin, recruits the Nup107-160 complex and thereby establishes a “seed” for new NPCs (Rasala et al., 2006; Franz et al., 2007). Next, the NPC seed comes into proximity of ER cisternae that cover chromatin for nuclear envelope reassembly. ER cisternae are perforated with nanoholes (also called fenestrations), whose architecture is topologically equivalent to that of nuclear pores. ER nanoholes associate with the NPC seed leading to integration of the seed into the nascent nuclear membrane. (Schellhaus et al., 2016; Otsuka et al., 2018; Bilir et al., 2019). NPC biogenesis is completed by sequential addition of the remaining nucleoporins (Dultz et al., 2008; Otsuka and Ellenberg, 2018). This post-mitotic pathway and the mechanistically distinct interphase pathway each form around half of a cell’s NPCs (Doucet et al., 2010; Dultz et al., 2010; Otsuka et al., 2016; Otsuka and Ellenberg, 2018). Both pathways require the induction and stabilization of high membrane curvature to create the nuclear pore. The nucleoporins Nup133, Nup53 (also called Nup35), Nup153, and Pom33 sense and potentially create membrane curvature. However, these properties are only essential for NPC assembly during interphase or in *S. cerevisiae* where the nuclear envelope remains intact throughout mitosis (Chadrin et al., 2010; Doucet et al., 2010; Vollmer et al., 2012, 2015). Additionally, the membrane-deforming proteins Reticulon4 and Tts1 contribute to NPC assembly in *S. cerevisiae* and interphase assembly in *X. laevis* (Dawson et al., 2009; Zhang and Oliferenko, 2014). For post-mitotic assembly, however, factors that control high-curvature ER formation and dynamics and that coordinate ER remodeling spatially and temporally with formation of the NPC seed are unknown.

Membrane-shaping within the cytoplasmic (also called peripheral) ER depends on reticulon homology domain (RHD)-proteins, Reticulon1-4 and REEP1-6 in mammals. RHD proteins bend membranes and thereby establish ER tubules and the curved edges of ER cisternae (Voeltz et al., 2006; Westrate et al., 2015; Shibata et al., 2009; Shibata et al., 2010). Additionally, RHD proteins are required for the formation of nanoholes within ER cisternae and localize to the edges of nanoholes (Schroeder et al., 2019). RHD proteins are enriched in these regions of high membrane curvature while low-curvature ER, in particular the nuclear envelope, was initially found to exclude RHD proteins (Voeltz et al., 2006; Shibata et al., 2010). The RHD protein REEP4, however, associates with the nuclear rim in addition to its expected localization to the peripheral ER (Schlaitz et al., 2013). Consistent with this observation, a proteomics study detected REEP4 as significantly enriched in the nuclear fraction compared to other RHD proteins (Itzhak et al., 2016). REEP4 and the closely related REEP3 position and shape the ER during mitosis (Schlaitz et al., 2013; Kumar et al., 2019). The unusual localization of REEP4 to the nuclear rim prompted us to investigate how it targets to the nuclear envelope and whether it plays a role in NPC formation.

In this study, we identify ELYS as a determinant of REEP4 targeting to the inner nuclear membrane (INM). We show that REEP4 is required for normal NPC levels and in particular for the formation of post-mitotically assembled NPCs, suggesting that REEP4 may coordinate high curvature ER with the ELYS-based NPC seed to promote NPC biogenesis during late mitosis.

## Results

### A pool of REEP4 localizes to the inner nuclear membrane

Endogenous REEP4 localizes to the cytoplasmic ER but also the nuclear envelope, in striking contrast to other RHD proteins including REEP5 and Reticulon4 (Schlaitz et al., 2013; Voeltz et al., 2006; Shibata et al., 2010). A C-terminally HA-tagged version of REEP4 similarly associates with the nuclear envelope (Figure 1A). We showed that REEP4-HA can functionally substitute for endogenous REEP4 during mitosis, is expressed at similar levels as the endogenous protein and does not cause ER morphology defects in transfected cells (Schlaitz et al., 2013; Kumar et al., 2019). To facilitate analysis of REEP4 in imaging assays we used REEP4-HA or related constructs that behaved equivalently.

**Figure 1.**
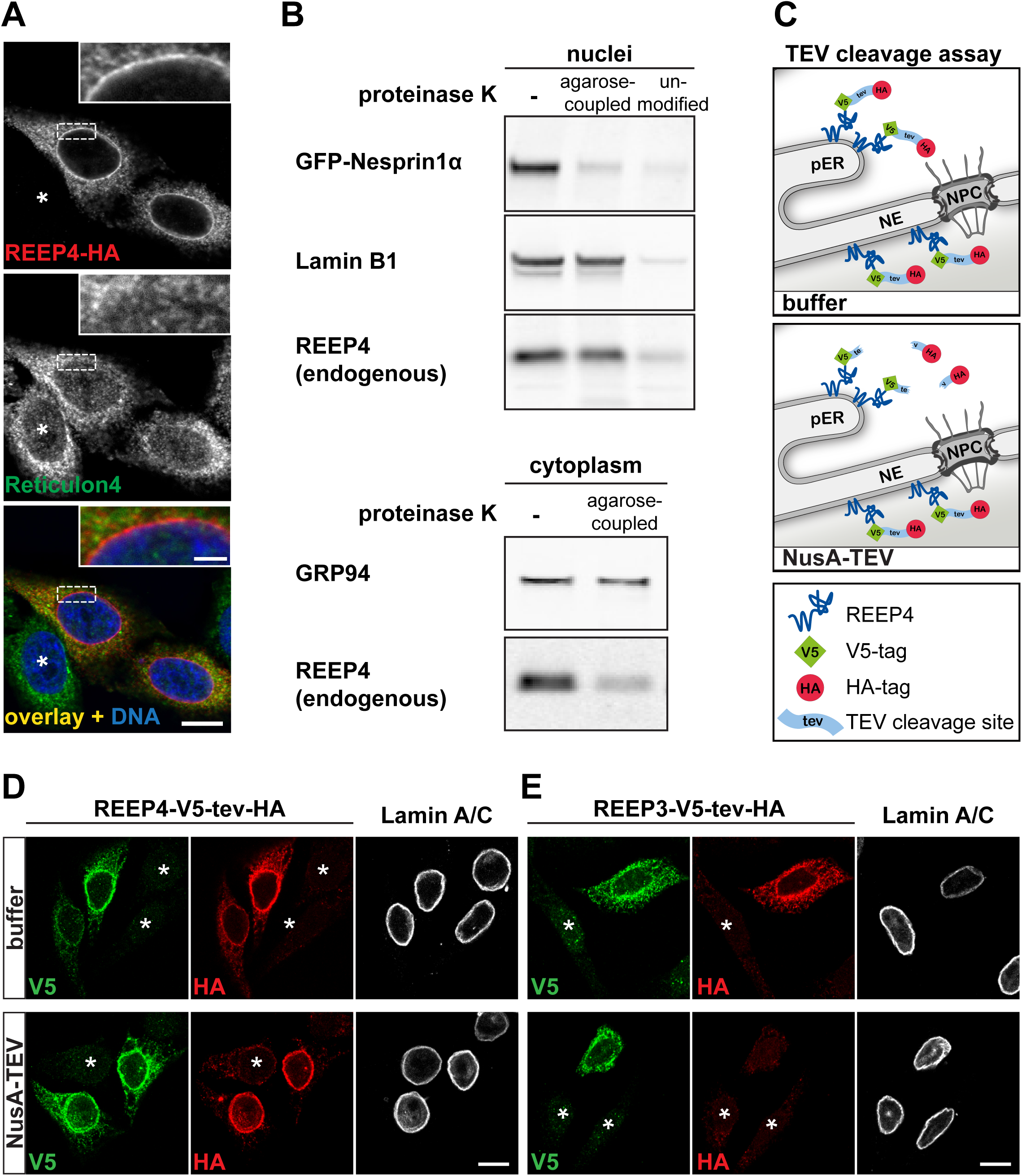
A pool of REEP4 localizes to the inner nuclear membrane. (A) HeLa cells expressing REEP4-HA, immunolabeled for the HA-tag and the endogenous RHD protein Reticulon4. DNA is stained with Hoechst dye. Inset shows part of the nucleus, scale bar is 2 µm. (B) HEK293T cells expressing the ONM marker GFP-Nesprin were fractionated into nuclei and cytoplasm. Upper panel: Nuclei were treated with agarose-coupled or unmodified proteinase K and analyzed by immunoblotting. Endogenous REEP4 in the nuclear fraction was largely resistant to agarose-proteinase K-treatment, similar to the INM protein Lamin B1. Lower panel: In the cytoplasmic fraction, agarose-coupled proteinase K degraded REEP4 but not the ER lumenal protein GRP94, suggesting that ER maintained its integrity. (C) Principle of the NusA-TEV cleavage assay. Semi-permeabilized cells expressing REEP4-V5-tev-HA are incubated with buffer or purified NusA-TEV. NusA-TEV removes cytoplasmic HA-tags. Remaining V5 marks REEP4-V5-tev-HA in expressing cells. HA-tags within the nucleus are protected from NusA-TEV cleavage. pER: peripheral (cytoplasmic) ER, NPC: nuclear pore complex, NE: nuclear envelope. (D), (E) NusA-TEV cleavage assays were performed on HeLa cells expressing (D) REEP4-V5-tev-HA or (E) REEP3-V5-tev-HA. Cells were immunostained for the V5-tag, HA-tag and Lamin A/C (for identification of the nuclear rim). (D) REEP4-HA staining persists at the nuclear rim. (E) REEP3-HA staining is abolished by NusA-TEV treatment suggesting that REEP3 is exclusively cytoplasmic. (A), (D), (E) Scale bars are 10 µm. Asterisks indicate untransfected cells.

We first asked whether REEP4 associates with the cytoplasmic face of the nuclear envelope (outer nuclear membrane, ONM) or the nucleoplasmic face of the nuclear envelope (inner nuclear membrane, INM). REEP4 is anchored in the ER through its N-terminal RHD and the soluble part of the protein is exposed to the cytoplasm or, possibly, the nucleoplasm (Park et al., 2010; Brady et al., 2015; Kumar et al., 2019). These properties allowed us to probe the topology of nuclear REEP4 using limited proteolysis. First, we isolated HEK293T cell nuclei and incubated them with either agarose-coupled or unmodified proteinase K. Agarose-proteinase K cannot enter an intact nucleus due to its large size and only degrades proteins of the ONM. The smaller unmodified proteinase K freely diffuses through nuclear pores and digests proteins of the ONM and INM. As expected, both proteinase K variants degraded the ONM protein Nesprin1α whereas the INM protein Lamin B1 was degraded by the free protease but not by agarose-proteinase K (Figure 1B, top panel). Endogenous REEP4 within the nuclear fraction persisted after agarose-proteinase K treatment, similar to Lamin B1. Importantly, agarose-proteinase K degraded REEP4 in the cytoplasmic fraction, confirming that REEP4 is in principle susceptible to agarose-proteinase K digest (Figure 1B, bottom panel). Second, we tagged REEP4 C-terminally with both, V5- and HA-tags, separated by a TEV protease cleavage site. Cells expressing REEP4-V5-tev-HA were first treated with digitonin to selectively permeabilize the plasma membrane while leaving organelle membranes intact (Adam et al., 1990) and then incubated with purified NusA-TEV protease, which is restricted to the cytoplasm (Theerthagiri et al., 2010; Ungricht et al., 2015). NusA-TEV should cleave the HA-tag from cytoplasmic REEP4-V5-tev-HA whereas intranuclear HA-tags should persist (Figure 1C). Indeed, after NusA-TEV-treatment, the cytoplasmic HA signal disappeared and only the nuclear rim remained HA-labeled, indicating that REEP4 was protected from TEV cleavage within the nucleus (Figure 1D). The closely related REEP3 could not be detected at the nuclear rim with this assay (Figure 1E) consistent with the observation that REEP3 is less abundant in nuclei compared to REEP4 (Itzhak et al., 2016). Thus REEP4 is present in a nuclear domain that is inaccessible to cytoplasmically-restricted proteases. Given that REEP4 is a membrane-associated protein and localizes to the nuclear rim, this domain should be part of the nuclear envelope and could be the general INM or the nuclear pore membrane, which is characterized by high curvature. In the following, we will subsume both these membrane regions under the term INM.

Our results demonstrate that a pool of REEP4 localizes to the INM. Our results, together with previous microscopy and proteomics studies (Voeltz et al., 2006; Shibata et al., 2010; Schlaitz et al., 2013; Itzhak et al., 2016), indicate that among RHD-containing proteins this INM localization is unique to REEP4.

### BioID identifies the nucleoporin ELYS as proximal to REEP4

To begin to elucidate targeting mechanisms and possible functions of REEP4 in the nucleus, we characterized its molecular environment using proximity-dependent biotin identification (BioID). In BioID, a biotin ligase-tagged protein of interest biotinylates proximal proteins, which are isolated by affinity capture and identified by mass spectrometry (Roux et al., 2012).

A fusion protein consisting of REEP4 and the engineered biotin ligase TurboID (Branon et al., 2018) localized to cytoplasmic ER and INM, as expected (Figure 2A). To control for non-specifically biotinylated proteins in the nucleus and cytoplasm, we performed BioID with TurboID fused to either a nuclear localization signal (NLS) or to REEP5 (Figure 2B). REEP4-TurboID, REEP5-Turbo-ID and TurboID-NLS were expressed, localized as expected and biotinylated nearby proteins (Figure S1). We performed three independent purifications and calculated the fold enrichment of proteins obtained with REEP4-TurboID versus the controls REEP5-Turbo-ID and TurboID-NLS (Mellacheruvu et al. 2013). The resulting list of REEP4-specific proximal proteins contained the known interaction partners Rab3GAP1, Rab3GAP2 and 14-3-3 proteins (Tinti et al., 2012) as well as REEP4 itself, validating our approach (Figure 2C). Remarkably, the most strongly enriched REEP4 proximal protein was the nucleoporin ELYS.

**Figure 2.**
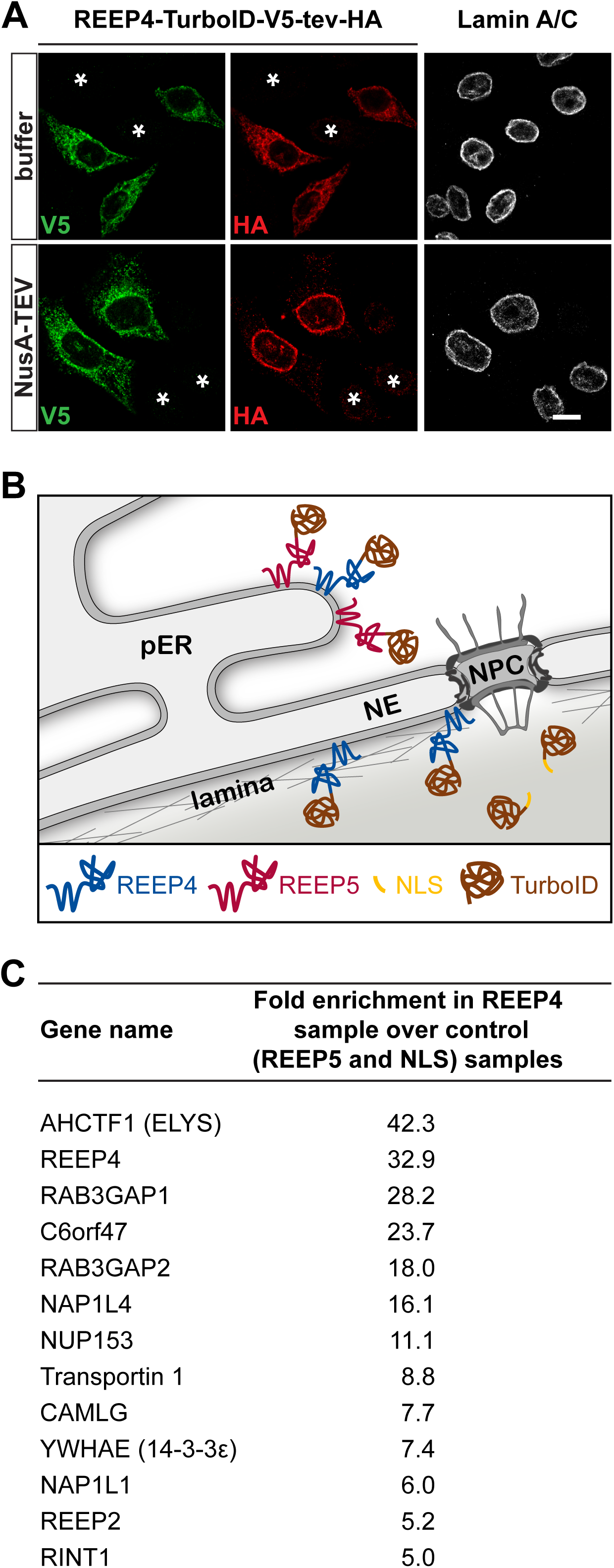
BioID experiment identifies the nucleoporin ELYS as proximal to REEP4. (A) NusA-TEV cleavage assay on HeLa cells expressing REEP4-TurboID-V5-tev-HA. Scale bar 10 µm. Asterisks indicate untransfected cells. (B) Rationale of the BioID approach. TurboID fusions of REEP4, REEP5 or a nuclear localization sequence (NLS) are expressed in HEK293 cells. REEP5-TurboID and TurboID-NLS serve to identify non-specifically biotinylated proteins in the peripheral ER and nucleoplasm, respectively. pER: peripheral ER, NE: nuclear envelope, NPC: nuclear pore complex. For localization and activity of all BioID constructs see figure S1. (C) Top hits for proteins identified as proximal to REEP4 with the BioID experiment. For each protein, fold enrichment in the REEP4-TurboID sample compared to the control samples REEP5-TurboID and TurboID-NLS is shown Hits five-fold or more enriched in the REEP4 sample are displayed.

### REEP4 and ELYS partially co-localize and ELYS promotes REEP4 INM targeting

To corroborate the BioID results, we first compared REEP4 and ELYS localization patterns by STED microscopy. REEP4 and ELYS appeared as partially overlapping puncta along the nuclear envelope, consistent with association of both proteins with nuclear pores (Figure 3A, B). In contrast, Lamin B1 showed a smooth distribution along the nuclear rim (Figure 3C).

**Figure 3.**
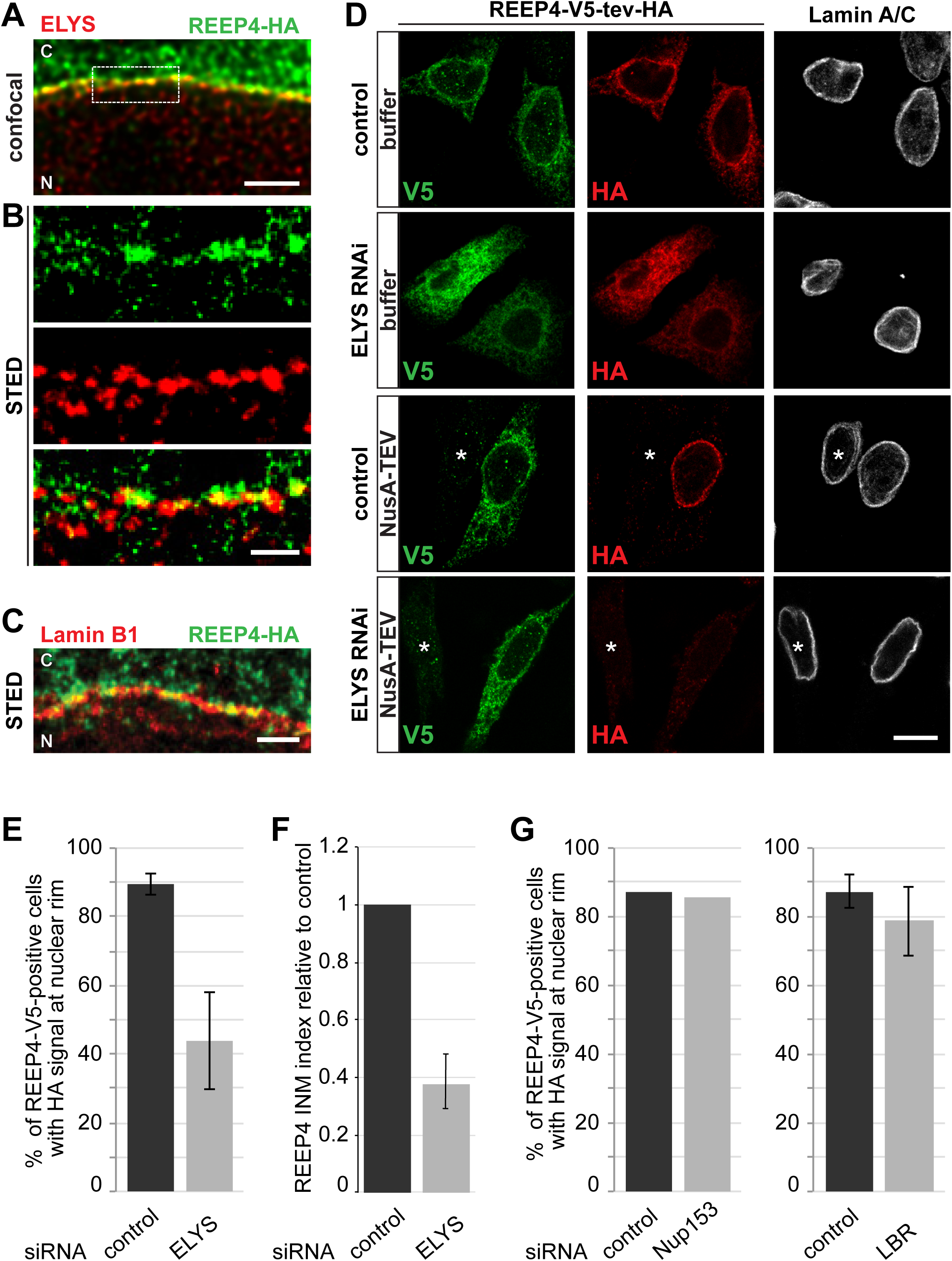
REEP4 and ELYS partially co-localize at the INM and ELYS promotes REEP4 INM targeting. (A) HeLa cell expressing REEP4-HA, immunolabeled for the HA-tag and ELYS and imaged by confocal microscopy. Part of the nuclear rim was acquired and the resulting image deconvolved. Scale bar is 2 µm. (B) Magnification of the region outlined in the confocal image in (A) acquired by STED microscopy. Scale bar is 500 nm. (C) Cell expressing REEP4-HA, stained for HA and Lamin B1 and imaged by STED microscopy. Scale bar is 1 µm. (A, C) C: cytoplasm, N: nucleoplasm. (D) NusA-TEV assay of HeLa cells expressing REEP4-V5-tev-HA, treated with control or ELYS siRNA. Cells were stained for V5-tag, HA-tag and Lamin A/C. Scale bar is 10 µm. Asterisks indicate untransfected cells. (E) Fraction of REEP4-V5-tev-HA-positive cells with HA-staining at the nuclear rim after NusA-TEV treatment in control and ELYS RNAi cells. Shown is the average of five experiments. (F) “REEP4 INM index” was determined by dividing HA signal intensity by V5 signal intensity for each cell. Mean value for REEP4 RNAi was normalized to the control. Shown is the average of five experiments with 25 cells analyzed per condition. (G) Fraction of REEP4-V5-tev-HA-expressing, NusA-TEV-treated cells with HA-staining at the nuclear rim in control cells or after depletion of Nup153 (average of two experiments with very similar outcomes is shown) or LBR (n=3). (E, G) At least 35 cells were analyzed per condition in a blinded manner. (E, F, G) Error bars are standard error of the mean (SEM).

The accumulation of membrane proteins at the INM depends on retention partners (Ungricht et al., 2015; Boni et al., 2015). We used the TEV cleavage assay to determine whether ELYS could be a retention partner for REEP4. Indeed, REEP4 targeting to the INM was strongly dependent on ELYS: REEP4 was present at the INM in 89% of control but only 44% of ELYS-depleted cells (Figure 3D, E). We also calculated the ratio of HA-signal intensity (representing the REEP4 nuclear pool) over V5-signal intensity (representing the entire REEP4 pool) to estimate how the fraction of nuclear REEP4 changed. This ratio, which we term REEP4 INM index, dropped by 60% after ELYS RNAi (Figure 3F). ELYS RNAi did not change expression levels of REEP4 (Figure S2A, S2B). Depletion of the nucleoporin Nup153, which was also identified as proximal to REEP4 (Figure 2C), did not diminish the fraction of cells with REEP4 at the INM, indicating that this phenotype is specific to ELYS RNAi (Figure 3G, S2A, S2C). The INM markers SUN2, Emerin, Lamin A/C and Lap2β were not mis-localized in ELYS RNAi (Figure S2E, F), suggesting that ELYS has no general role in retention of INM proteins. However, the INM protein LBR was mis-targeted after ELYS RNAi as reported previously (Figure S2E, far right column; Mimura et al., 2016). Yet, LBR depletion did not affect INM targeting of REEP4 (Figure 3G, S2A, S2D), ruling out that REEP4 mis-localization resulted from disturbed LBR distribution. Finally, HA-positive rim labeling of the independent INM marker HA-tev-V5-Lap2β was not reduced by ELYS RNAi, suggesting that depletion of ELYS did not cause NusA-TEV protease to aberrantly enter nuclei (Figure S2F).

These results indicate that ELYS directly contributes to the targeting of REEP4 to the INM.

### REEP4 is required for normal NPC levels at the nuclear envelope

The NPC biogenesis factor ELYS targets REEP4 to the INM, hinting at a role for REEP4 in NPC assembly. We analyzed NPC levels using three different markers: ELYS, FG-nucleoporins detected by the Mab414 antibody, and RanBP2 (Figure 4A, B). All three markers were decreased by about 20% after REEP4 RNAi while Lamin B1 was not diminished. NPC reduction was observed with three distinct REEP4 siRNAs (Figure S3A, B). However, REEP4 depletion did not affect ELYS protein levels (Figure S3C).

**Figure 4.**
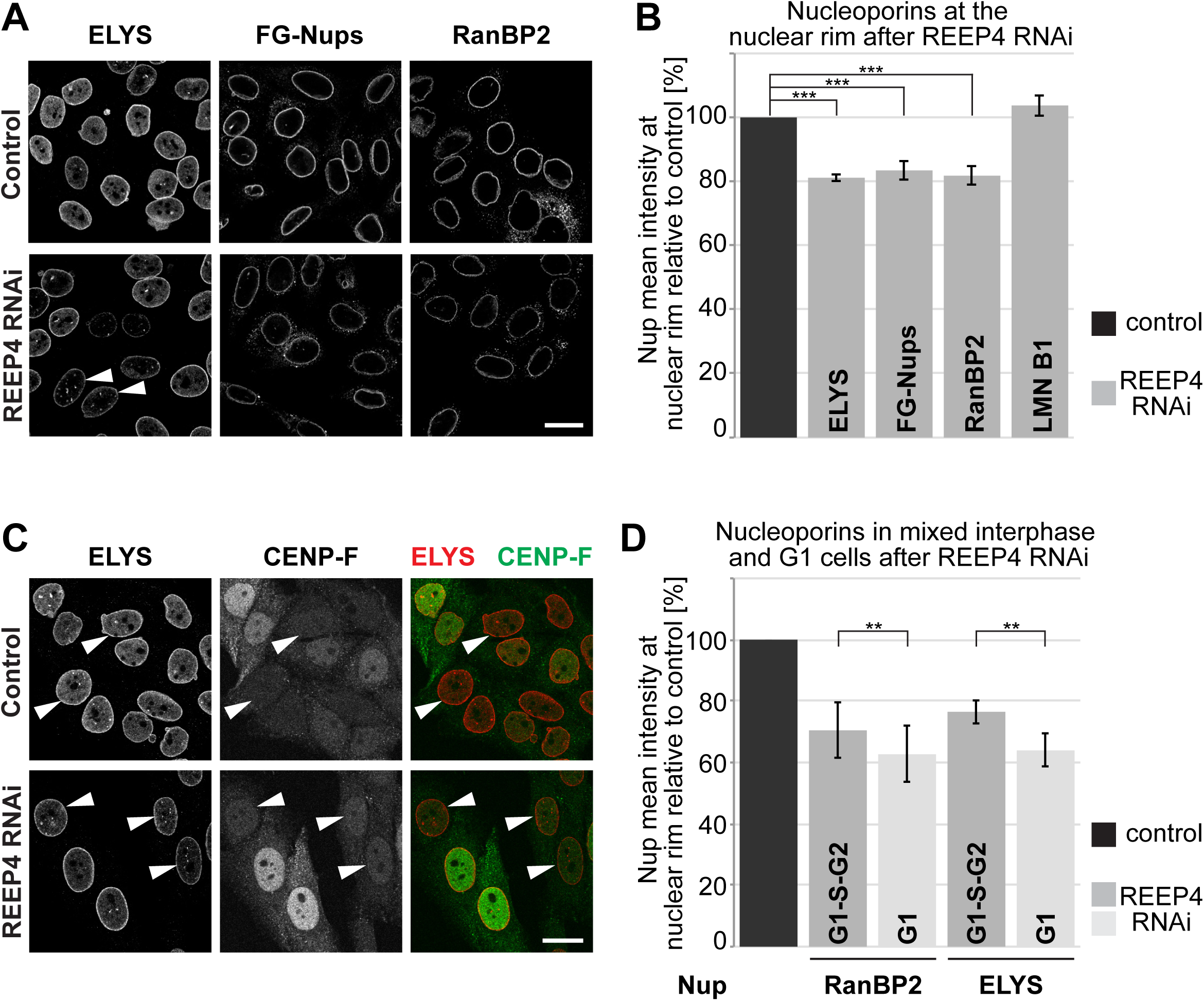
REEP4 is required for normal NPC densities. HeLa cells were treated with non-targeting control siRNA or REEP4 siRNA, fixed, immunolabeled for indicated nucleoporins and (C) CENP-F, and imaged by confocal microscopy. (A, C) Representative images. Scale bars are 20 µm. (A) Arrowheads indicate cells with an increased number of nucleoplasmic ELYS foci after REEP4 RNAi. For quantification see figure S3D. (B, D) Mean intensities for the respective NPC markers at the nuclear rim were measured in control and REEP4 RNAi cells and normalized to control conditions. (B) At least 100 cells were analyzed per condition, shown are means from three (LMNB1), four (ELYS, RANBP2) or five (FG-Nups) experiments, error bars are SEM. For each single experiment, nucleoporin levels in REEP4 RNAi were significantly different from controls with p-values smaller than 0.001. (C) Arrowheads indicate cells with low CENP-F signal, which are in G1 phase. (D) Mean nucleoporin intensities were measured in either the entire interphase population (G1-S-G2) or in G1 cells, identified by lack of CENP-F staining. Nucleoporin levels shown are relative to the respective control (either total interphase or G1). Averages from three (RanBP2) or five (ELYS) experiments are shown with at least 100 cells analyzed for the total interphase population and 24 cells for G1. In each single REEP4 RNAi data set, G1 means were significantly different from total interphase means with p values <0.01. (B, D) Error bars are SEM. Welch’s t-test was used to calculate p-values.

ELYS localizes to NPCs but additionally to nucleoplasmic structures called GLFG bodies (Morchoisne-Bolhy et al., 2015). The abundance of ELYS-positive nuclear bodies increased upon REEP4 depletion, suggesting that ELYS accumulates in these structures if it is not incorporated into NPCs (Figure 4A, S3D). Exogenous expression of REEP4-HA increased NPC density in wild-type cells and partially rescued NPC levels in REEP4 RNAi cells (Figure S3E, F).

REEP4 shapes the ER during mitosis and ELYS initiates post-mitotic NPC assembly, suggesting that both proteins might collaborate in NPC biogenesis in late mitosis. We used the cell cycle-regulated protein CENP-F (Liao et al., 1995) to identify early interphase (G1) cells because during G1, NPC levels are dominated by the contribution of post-mitotic assembly. To determine whether REEP4 contributes mainly to post-mitotic NPC assembly, we analyzed the nucleoporins ELYS and RanBP2 in mixed interphase and G1 cells. For both, ELYS and RanBP2, REEP4 RNAi resulted in a significantly larger reduction in NPC densities in G1 compared with mixed interphase cells (Figure 4C, D).

These results demonstrate that REEP4 is required for normal levels of NPCs within the nuclear envelope and indicate that REEP4 acts predominantly in post-mitotic NPC assembly.

## Discussion

We report here that REEP4, a protein creating and residing in high curvature ER, is recruited to the INM by the NPC biogenesis factor ELYS. ELYS and REEP4 are close to each other in the cell and may interact directly, and both proteins promote NPC formation during late mitosis. Based on these observations, we speculate that REEP4 aids the association of high curvature ER, possibly nanoholes within ER cisternae, with chromatin-bound ELYS during early stages of post-mitotic NPC formation to enhance the efficiency of the assembly process. Depletion of ELYS leads to an approximately 40% reduction in NPC densities (Jevtic et al., 2019) while REEP4 RNAi leads to a 20% reduction, in agreement with our model that REEP4 supports but is not essential for NPC assembly. REEP4-HA restores normal high curvature morphology of the ER during metaphase in REEP3/4 RNAi cells (Kumar et al., 2019) but did not fully rescue the REEP4 RNAi-mediated defect in NPC levels, possibly because the amounts of expressed REEP4-HA are insufficient or because REEP4-HA interacts less efficiently with ELYS than endogenous REEP4.

A direct interaction between REEP4 and ELYS in anaphase could mediate the localization of REEP4 to the forming nucleoplasmic face of the nuclear envelope. This post-mitotic nuclear inclusion of REEP4 would obviate a need for transport of REEP4 through NPCs. REEP4 may alternatively or additionally target to the nuclear membrane throughout interphase by passive diffusion through peripheral channels of the NPC and retention at the INM (Ungricht et al., 2015). ELYS depletion could in theory change the properties of NPCs, e.g. by restricting the width of the peripheral channels, and thereby indirectly lead to the exclusion of INM protein with large cytoplasmic domains. However, the cytoplasmic domain of REEP4 with 190 amino acids is smaller than that of the INM proteins we studied and found to localize normally (cytoplasmic domains of Emerin, SUN2 and Lap2β are 221, 212 and 410 amino acids, respectively) and it is therefore unlikely that REEP4 becomes excluded due to a secondary structural defect in the nuclear pore.

Atlastins, a different class of ER shaping proteins, are also required for NPC biogenesis by maintaining proper structure of peripheral ER, which ensures normal mobility of INM proteins within the ER network (Pawar et al., 2017). After REEP3/4 RNAi, interphase ER maintains a normal morphology and REEP4 depletion alone does not affect mitotic ER morphology (Schlaitz et al., 2013; Kumar et al., 2019). REEP4 is furthermore not required for accumulation of the membrane-associated Lamin B1 at the INM (Figure 4B). Thus, it is improbable that REEP4 RNAi reduces NPC levels due to disruption of ER morphology.

REEP4 is present at the INM in nearly 90% of control cells, meaning that REEP4 remains at the INM after post-mitotic NPC formation. REEP4 might be stored within the nucleus to promote the conversion of ER membrane to higher curvature in the next mitosis (Puhka et al., 2012; Kumar et al., 2019).

In summary, REEP4 is the first RHD protein observed at the INM, and it is required for normal incorporation of NPCs into the nuclear membrane. REEP4 contributes primarily to post-mitotic NPC formation, possibly by linking high-curvature ER and NPC precursors during anaphase. How REEP4 interacts with membranes and ELYS to promote NPC formation are important topics for future research.

## Materials and Methods

### Cell culture, siRNAs, transfections, generation of REEP4 knockout line

HeLa cells (ATCC CCL-2, authenticated by UC Berkeley tissue culture facility) and HEK293T cells (ATCC CRL-11268) were cultured in DMEM, supplemented with 10% fetal bovine serum (FBS), at 37°C in a humidified 5% CO_2_ incubator. Cells were checked for mycoplasma contamination approximately every six months and were always negative. For plasmid transfections or siRNA/plasmid co-transfections, 1.8×10^5^ cells were seeded in one well of a 24-well plate and transfected the next day with 0.4 µg plasmid DNA using Lipofectamine 2000 (Invitrogen) according to the manufacturer’s instructions. Four to six hours after transfection, cells were split onto coverslips and incubated for a total of 72 hours with siRNAs/plasmids before fixation. siRNAs were transfected concomitant with cell seeding using Lipofectamine RNAiMax (Invitrogen) according to the manufacturer’s instructions, the cells were split onto coverslips the next day and incubated for a total of 72 hours with siRNAs before fixation. siRNAs used were from ThermoFisher with the following IDs: Negative control siRNA: AM4611, REEP4 siRNA#1: #32438, REEP4 siRNA#2: s37272, REEP4 siRNA#3: s37270, ELYS siRNA: #108720, LBR siRNA: s8101, Nup153 siRNA: s19375.

### TEV cleavage assay

Cells were subjected to TEV assays 48 hours after transient transfection of the respective TEV assay reporters. Cell were semi-permeabilized by a 10-minute incubation on ice with freshly prepared permeabilization buffer (20 mM HEPES pH 7.5, 110 mM CH3COOK, 5 mM (CH3COO)2Mg, 250 mM sucrose, 0.5 mM EGTA) containing 0.0025% digitonin (from 5% stocks in water, prepared the same day), washed three times with permeabilization buffer without digitonin and incubated with purified NusA-TEV protease (Theertagiri et al., 2010) for 15 minutes at 30°C. Immediately following incubation with NusA-TEV, coverslips were fixed with 4% formaldehyde in PBS and processed for immunofluorescence. The TEV assay procedure was adapted from Ungricht et al., 2015. Purified NusA-TEV protease was kindly provided by Rosemarie Ungricht and Ulrike Kutay.

### Plasmids used in this study

Plasmids encoding human REEP3-HA and REEP4-HA have been described previously (Schlaitz et al., 2013; Kumar et al., 2019). The V5-tag and TEV cleavage site sequences were inserted into these plasmids based on Gibson assembly (NEB) using an oligo encoding for these features. For generating HA-tev-V5-Lap2β, pEGFP-Lap2β was used as the host plasmid (obtained from Euroscarf; Beaudouin et al., 2002). The plasmid encoding the TurboID enzyme as well as Turbo-ID-NLS were obtained from Addgene (Branon et al., 2018). REEP4-TurboID-HA and REEP5-TurboID-HA constructs for the generation of inducible cell lines were generated by combining the sequences of REEP4 or REEP5 (from HA-REEP5, Schlaitz et al., 2013) with the Turbo-ID-HA-tag in pcDNA5/FRT/TO vectors using Gibson assembly. GFP-Nesprin1α was a gift from Andreas Merdes (Espigat-Georger et al., 2016).

### Immunofluorescence and light microscopy

Primary and secondary antibodies used are specified in the tables below. DNA was labeled with Hoechst 33342 (Merck). Coverslips were mounted with ProLong Diamond (ThermoFisher).

For immunofluorescence following TEV assays, cells grown on coverslips were permeabilized with 0.1% Triton X-100 (Merck) in PBS and blocked with 5% normal donkey serum (Abcam) in PBS. Primary antibodies were diluted in PBS/0.1% Tween.

For immunofluorescence of nucleoporins, cells grown on coverslips were fixed with 4% formaldehyde/PBS, then blocked and permeabilized by incubation with immunofluorescence buffer (1% BSA, 0.02% SDS, 0.1% Triton X-100 in PBS). All antibody incubation and wash steps were performed using this buffer (Gomez-Cavazos and Hetzer, 2015). Primary antibody incubations with anti-NPC antibodies were performed over night at 4°C.

STED imaging was performed on a Leica TCS SP8 3X STED system (Leica Microsystems) HC PLAPO 93×/1.30 numerical aperture (NA) glycerol objective lens or a HC PLAPO 100×/1.40 NA STED oil objective lens. STED images were acquired by using a white light laser at 635 nm and a STED laser of 775 nm. HyD detectors (Leica Microsystems) were used for signal detection of STED samples. For confocal imaging either a HCX PLAPO 63×/1.40 NA oil objective lens on the TCS SP8 system or a 63×/1.40 NA Plan-Apochromat oil objective lens on a LSM 780 system (Zeiss) was used.

### Primary antibodies used

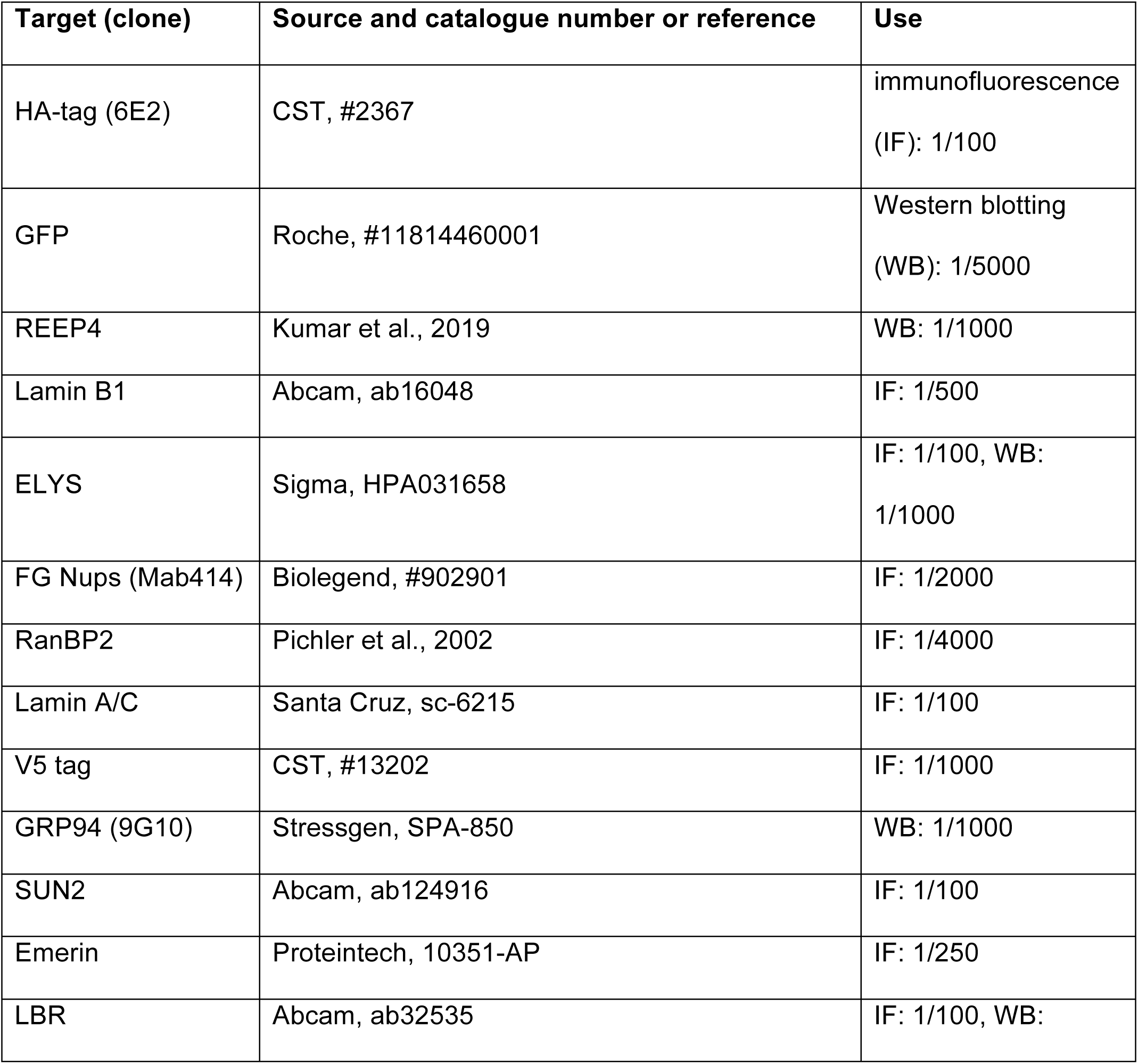
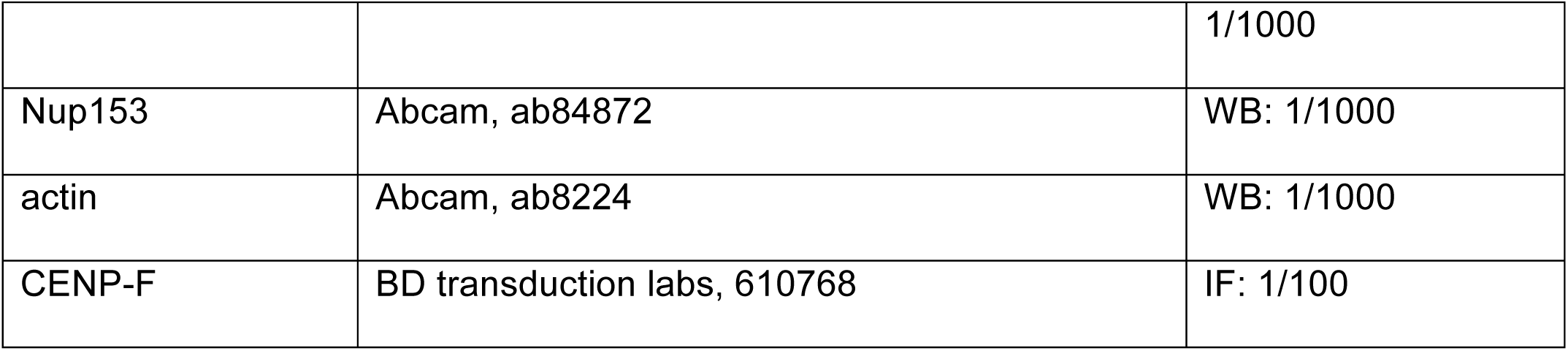

| Target (clone) | Source and catalogue number or reference | Use |
| --- | --- | --- |
| HA-tag (6E2) | CST, #2367 | immunofluorescence<br>(IF): 1/100 |
| GFP | Roche, #11814460001 | Western blotting<br>(WB): 1/5000 |
| REEP4 | Kumar et al., 2019 | WB: 1/1000 |
| Lamin B1 | Abcam, ab16048 | IF: 1/500 |
| ELYS | Sigma, HPA031658 | IF: 1/100, WB:<br>1/1000 |
| FG Nups (Mab414) | Biolegend, #902901 | IF: 1/2000 |
| RanBP2 | Pichler et al., 2002 | IF: 1/4000 |
| Lamin A/C | Santa Cruz, sc-6215 | IF: 1/100 |
| V5 tag | CST, #13202 | IF: 1/1000 |
| GRP94 (9G10) | Stressgen, SPA-850 | WB: 1/1000 |
| SUN2 | Abcam, ab124916 | IF: 1/100 |
| Emerin | Proteintech, 10351-AP | IF: 1/250 |
| LBR | Abcam, ab32535 | IF: 1/100, WB: |
|  |  | 1/1000 |
| Nup153 | Abcam, ab84872 | WB: 1/1000 |
| actin | Abcam, ab8224 | WB: 1/1000 |
| CENP-F | BD transduction labs, 610768 | IF: 1/100 |

### Secondary antibodies used

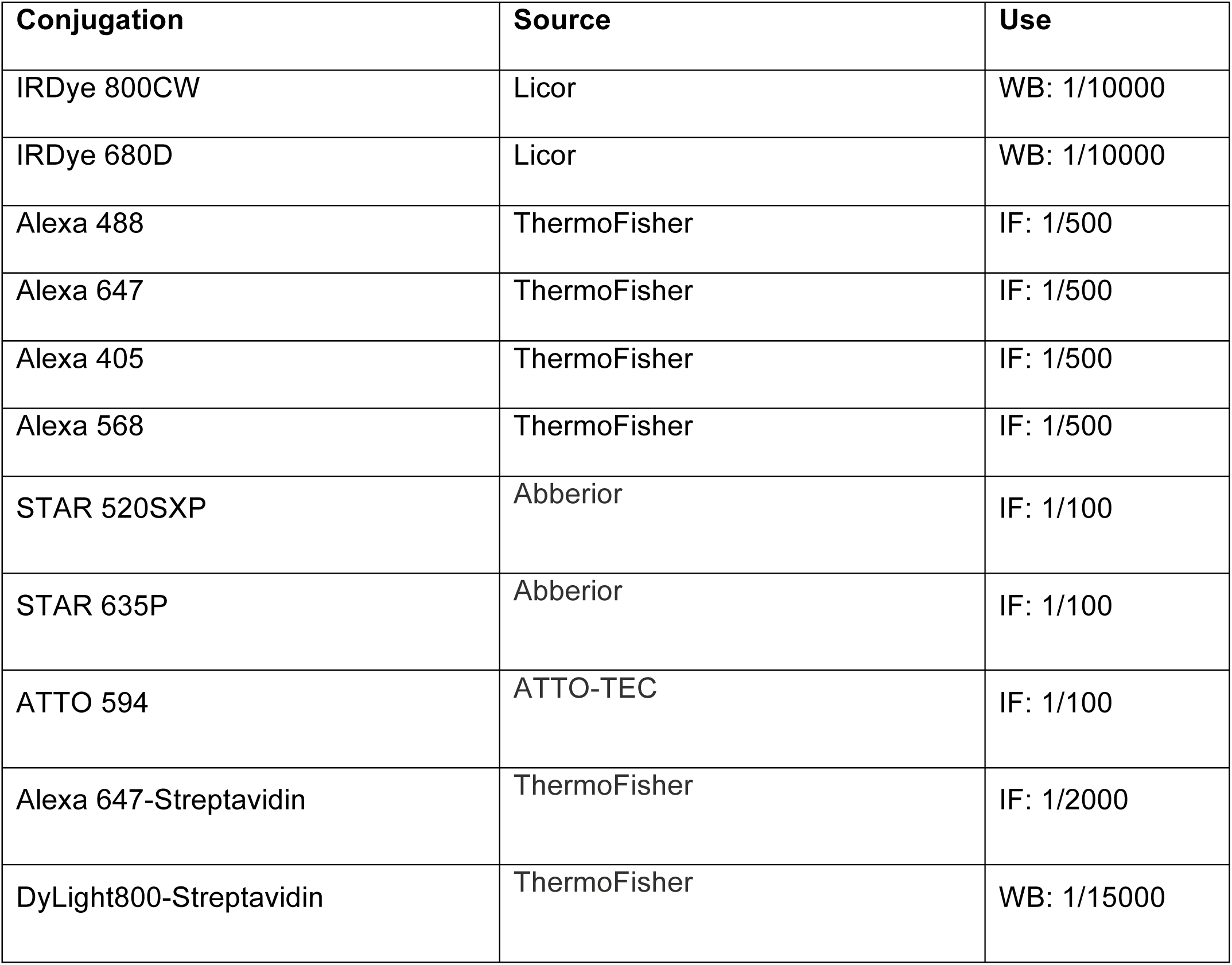

| **Conjugation** | **Source** | **Use** |
| --- | --- | --- |
| IRDye 800CW | Licor | WB: 1/10000 |
| IRDye 680D | Licor | WB: 1/10000 |
| Alexa 488 | ThermoFisher | IF: 1/500 |
| Alexa 647 | ThermoFisher | IF: 1/500 |
| Alexa 405 | ThermoFisher | IF: 1/500 |
| Alexa 568 | ThermoFisher | IF: 1/500 |
| STAR 520SXP | Abberior | IF: 1/100 |
| STAR 635P | Abberior | IF: 1/100 |
| ATTO 594 | ATTO-TEC | IF: 1/100 |
| Alexa 647-Streptavidin | ThermoFisher | IF: 1/2000 |
| DyLight800-Streptavidin | ThermoFisher | WB: 1/15000 |

Total protein on Western blots was detected with Revert700 Total Protein Stain (Licor).

### Image analysis and quantifications

Fiji was used for all microscopy image analysis (Schindelin et al., 2012). Microscopy images from NusA-TEV treated samples and of ELYS intranuclear clusters (Figures 3E and S3D) were analyzed and classified using the “Blind analysis Tools”-Plugin developed by Astha Jaiswal and Holger Lorenz (https://imagej.net/Blind_Analysis_Tools).

For quantification rim HA signal in the TEV assays for REEP4 INM index (Figure 3F) or nucleoporin densities (Figure 4B, D), background subtraction of the raw images was performed using a rolling ball algorithm with a diameter of 150 pixels (corresponding to the average cell size). Next, a mask was generated for the nuclear rim using either the staining of the nucleoporin itself or the staining of LaminA/C or LaminB1 in a separate channel. Segmentation was performed by applying a combination of the Gaussian filter, the ‘make Binary’ tool, iterative erosion and iterative dilation operations. The segmentation was corrected manually for ‘out of focus-regions’, and the final mask covered at least 30% of the whole rim. Mean Nup fluorescence intensities were measured in the resulting regions of interest. For quantification of CENP-F signal (Figure 4D), CENP-F mean intensity was measured inside the nuclear rim mask that was generated as described above. For quantification of V5 mean intensity for the REEP4 INM index, a mask was generated using the V5 signal.

### Proteinase K assay

HEK293T cells were lysed with a dounce homogenizer in S250 buffer (10 mM HEPES pH7.5, 250 mM sucrose, 50 mM KCl, 2.5 mM MgCl2) supplemented with 40 µM MG132 and 1 mM DTT, and spun for 10 minutes at 1,000xg to obtain cytoplasmic (supernatant) and crude nuclear (pellet) fractions. The fractions were incubated with buffer only, unmodified proteinase K (Sigma) or agarose-coupled proteinase K (Sigma) in lysis buffer for 30 minutes at room temperature. Reactions were stopped by addition of PMSF (Sigma). Samples were supplemented with Laemmli buffer and analyzed by SDS-PAGE and immunoblotting.

### Generation of stable cell lines for the inducible expression of TurboID constructs

For the generation of stable cell lines, Flp-In HEK293 cells (Gossen et al., 1995) were used and maintained in DMEM supplemented with 10% v/v FBS, 1x pyruvate, 1x GlutaMAX, 1% (v/v) penicillin/streptomycin at 37 °C under 5% CO_2_. 10 µg/ml blasticidin was added to select for cells that express the Tet repressor and 100 µg/ml zeocin to select for cells that have no gene of interest inserted downstream of the Tet promoter. Mycoplasma testing was performed before experiments and showed no contamination. Co-transfection of the Flp-Recombinase plasmid and the pcDNA5/FRT/TO vector containing the gene of interest (REEP4-TurboID-HA, REEP5-TurboID-HA or HA-TurboID-3xNLS) was performed in a 1:3 ratio in cells growing in the medium described above but without antibiotics. After 48 hours, selection of positive clones was started by changing the medium to DMEM supplemented with 10% v/v FBS, 1x pyruvate, 1x GlutaMAX, 1% (v/v) penicillin/streptomycin, 100 µg/ml hygromycin, 10 µg/ml blasticidin. Selection medium was changed every 2-4 days until colonies appeared after approximately 8-10 days. Individual colonies were isolated and grown to confluency in 24 well plate wells (with 50 µg/ml hygromycin). Clones were tested for successful insertion of the construct by inducing its expression with 100 ng/ml doxycycline for 24 hours.

### BioID experiment

Prior to large scale BioID experiments we determined concentrations of doxycycline for induction of all TurboID constructs to a level similar to endogenous REEP4. The doxycycline concentrations used were: REEP4-TurboID: 4 ng/ml, REEP5-TurboID: 0.35 ng/ml, TurboID-3xNLS: 2 ng/ml. 48 hours prior to harvest, cells were seeded in 15 cm dishes at a density of ∼20% and induced with doxycycline 24 hours prior to cell harvest. For biotinylation of proteins, biotin was added to a final concentration of 500 µM 1 hour prior to cell harvest. Cells were harvested by scraping, washed in PBS and snap frozen in liquid nitrogen.

Cells were lysed at room temperature by resuspension in 1 ml lysis buffer (50 mM Tris-HCl pH 7.5, 500 mM NaCl, 0.4% SDS, 5 mM EDTA, 1 mM DTT, 1 mM PMSF, Complete Protease Inhibitors (Roche), 40 µM MG132 (Sigma), PhosStop phosphatase inhibitors (Roche)) and passed ten times through a 20G needle. Subsequently, cell lysates were sonicated in a BioRuptur (Diagenode) for 10 intervals of 30 seconds high power, 30 seconds off to break protein aggregates and homogenize the lysate. 200 µl of 10% Triton X-100 were added to re-solubilize precipitated SDS and the samples were sonicated once more. 2.13 ml of 50 mM Tris pH 7.5 containing Complete Protease Inhibitors (Roche) and PhosStop phosphatase inhibitors (Roche) were added to reduce the salt concentration of the lysate to 150 mM, and lysates were distributed to two tubes. Lysates were cleared by centrifugation at 15,000xg for 10 min, 3 ml of supernatants were combined with 50 µl of equilibrated streptavidin coated beads (GE 17-5113-01) and incubated for 3 hours at 4°C. After incubation, the beads were pelleted at 2,500 xg for 2 min at 4°C and washed twice for 5 min each in a volume of 500 µl in the following buffers. Wash 1: Adjusted lysis buffer, 4°C. Wash 2: 2% SDS, room temperature. Wash 3: 50 mM HEPES pH 7.4, 1 mM EDTA, 500 mM NaCl, 1% Triton X-100, 0.1% Sodium deoxycholate, room temperature. Wash 4: 50 mM Tris-HCl pH 7.5, 50 mM NaCl, 0.1% Triton X-100 (procedure adapted from Schopp and Bethune, 2018)

To evaluate the success of the pulldown, 10% of beads were boiled at 95°C for 10 minutes with sample buffer containing 2 mM biotin and analyzed by immunoblotting. The remaining beads were washed once in urea buffer (8 M Urea, 100 mM Tris-HCl pH 8.8), resuspended in 50 µl urea buffer and stored at -80°C until mass spectrometry analysis.

### Mass spectrometry analysis of BioID samples

#### 1. On-Bead Tryptic Proteolysis

Streptavidin agarose beads were washed extensively with 8 M urea in 0.1 M Tris/HCl pH 8.5 followed by reduction and alkylation steps. Finally, the beads were washed with 50 mM ammonium bicarbonate and incubated with trypsin overnight in 50 mM ammonium bicarbonate at 37°C. The resulting peptides were collected via centrifugation at 1,000g for 5 min. The beads were then rinsed with 50 mM ammonium bicarbonate and this second tryptic fraction was pooled with the first one. After acidification, tryptic peptides were desalted by Stage Tipping using Empore C18 47mm disks, then dried in Speed-Vacuum. Further, they were resuspended in 5% formic acid and 5% acetonitrile for LC-MS/MS analysis.

#### 2. Data Acqusition

Peptides were analyzed by C18 nanoflow reversed-phase HPLC (Dionex Ultimate 3000 RSLC nano, Thermo Scientific) combined with orbitrap mass spectrometer (Q Exactive HF Orbitrap, Thermo Scientific). Samples were separated in an in-house packed 75 µm i.d. × 35 cm C18 column (Reprosil-Gold C18, 3 µm, 200 Å, Dr. Maisch) using 70 minute linear gradients from 5-25%, 25-40%, 40-95% acetonitrile in 0.1% formic acid with 300 nL/min flow in 90 minutes total run time. The scan sequence began with an MS1 spectrum (Orbitrap analysis; resolution 120,000; mass range 400–1,500 m/z; automatic gain control (AGC) target 1e6; maximum injection time 60 ms). Up to 20 of the most intense ions per cycle were fragmented and analyzed in the orbitrap with Data Dependent Acquisition (DDA). MS2 analysis consisted of collision-induced dissociation (higher-energy collisional dissociation (HCD)) (resolution 30,000; AGC 1e5; normalized collision energy (NCE) 27; maximum injection time 60 ms). The isolation window for MS/MS was 2.0 m/z.

#### 3. Data Processing

The data sets were searched against the human Swissprot/Uniprot database. Proteome Discoverer 1.4. software was used to identify proteins. Carbamidomethylation of cysteine was used as fixed modification and acetylation (protein N-termini) and oxidation of methionine were used as variable modifications. Maximal two missed cleavages were allowed for the tryptic peptides. The precursor mass tolerance was set to 10 ppm and both peptide and protein false discovery rates (FDRs) were set to 0.01. The other parameters were used with default settings. The database search was performed against the human Uniprot database (release 2016) with taxonomy Homo sapiens using Mascot and Sequest.

### Analysis of mass spectrometry data

Fold change calculation of identified proteins (Sample of interest / controls) was done using REPRINT (Resource for Evaluation of Protein Interaction Networks, Mellacheruvu et al. 2013, https://reprint-apms.org). In our final analysis, we considered proteins with a fold change equal to or larger than five, which were detected in at least two out of three replicates with an average of at least two peptide spectrum matches.

## Acknowledgements

We are indebted to Ulrike Kutay and Rosemarie Ungricht for advice on the TEV assay procedure, the generous gift of purified NusA-TEV protease and thoughtful discussions. We thank Frauke Melchior and her group for the kind gift of anti-RanBP2 antibody as well as discussions and advice throughout the project. We appreciate the kind gift of GFP-Nesprin1α plasmid from Andreas Merdes. We are grateful to Holger Lorenz, Astha Jaiswal and Christian Hörth (ZMBH) for expert light microscopy and image analysis support, and to members of the Schiebel, Schuck and Lemberg labs for discussions, reagents and advice. Many thanks to Rebecca Heald for comments on the manuscript.

This work was funded by the German Research Foundation (DFG, project no. SCHL1876/2-1, A.S.). A.S. is supported by a Short Term-Fellowship of the Chica and Heinz Schaller Foundation. B.G. is a graduate student of the HBIGS Graduate School of Molecular and Cellular Biology of Heidelberg University.

