## Supplemental Data for "REEP4 is recruited to the inner nuclear membrane by ELYS and promotes nuclear pore complex formation"

### Figure S1

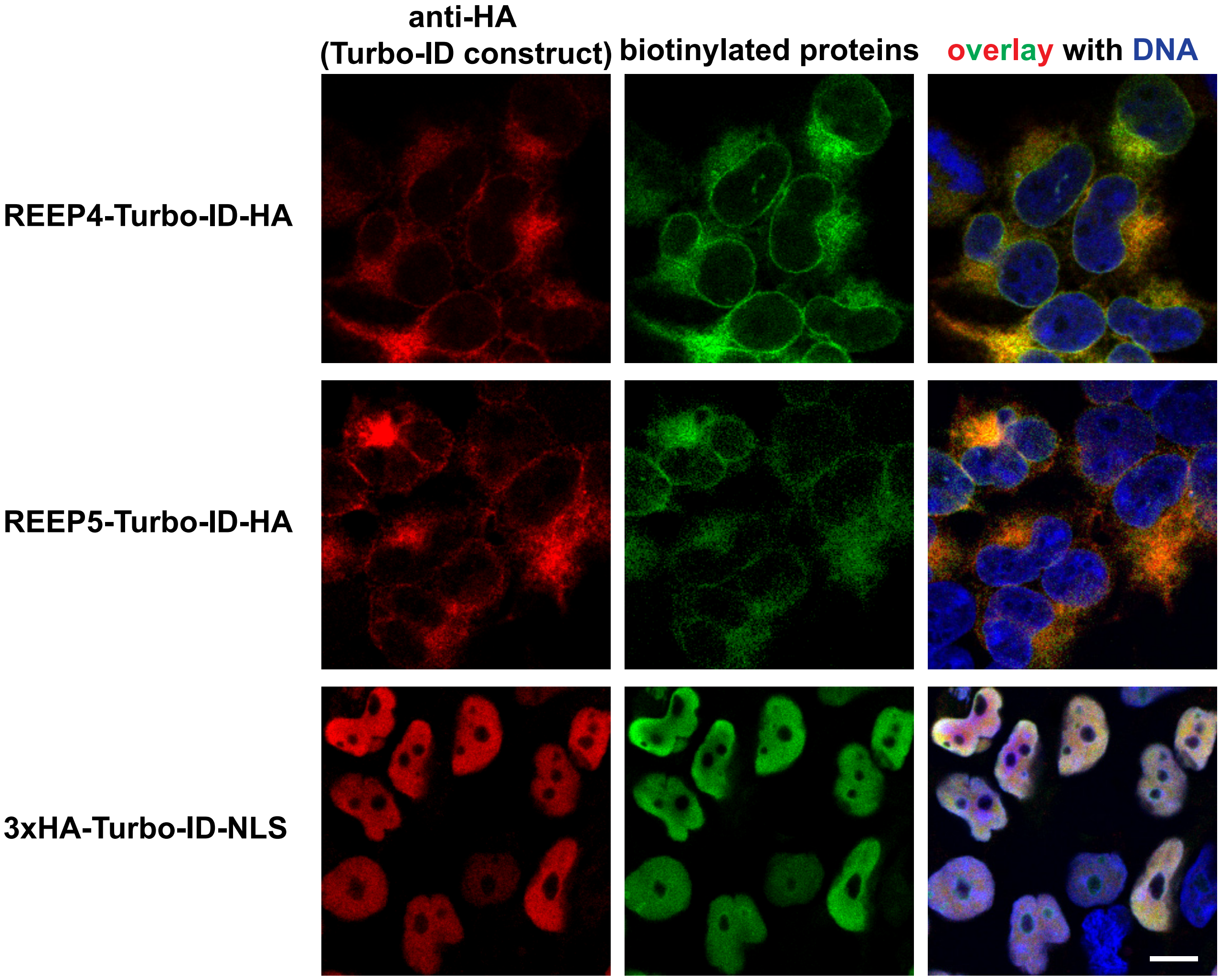
anti-HA
(Turbo-ID construct)
biotinylated proteins
overlay with DNA
REEP4-Turbo-ID-HA
REEP5-Turbo-ID-HA
3xHA-Turbo-ID-NLS

### Figure S2

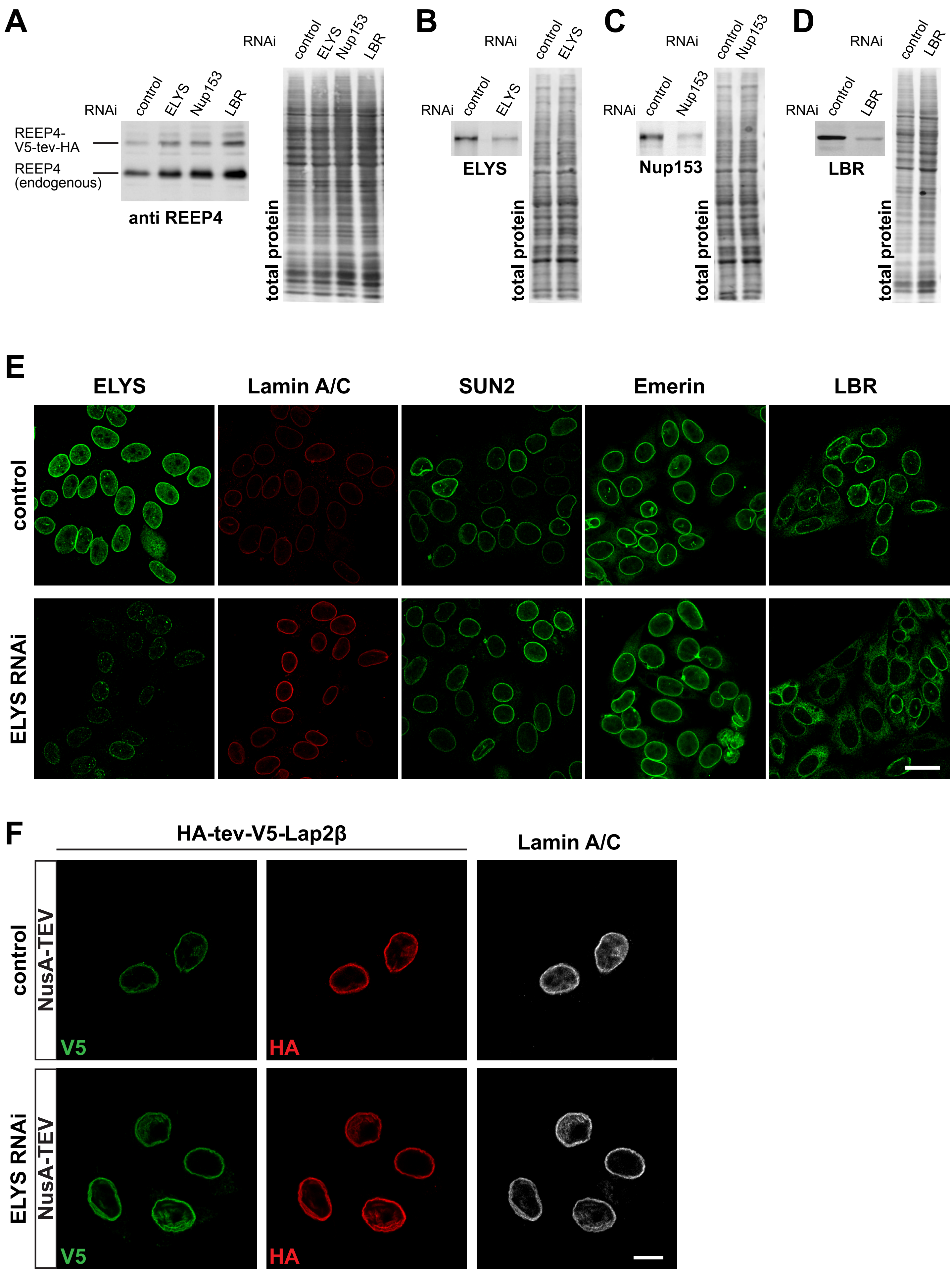
A
RNAi
control
ELYS
Nup153
LBR
REEP4-V5-tev-HA
REEP4 (endogenous)
anti REEP4
total protein
B
ELYS
C
Nup153
D
LBR
E
ELYS
Lamin A/C
SUN2
Emerin
LBR
control
ELYS RNAi
F
HA-tev-V5-Lap2β
Lamin A/C
NusA-TEV
V5
HA

### Figure S3

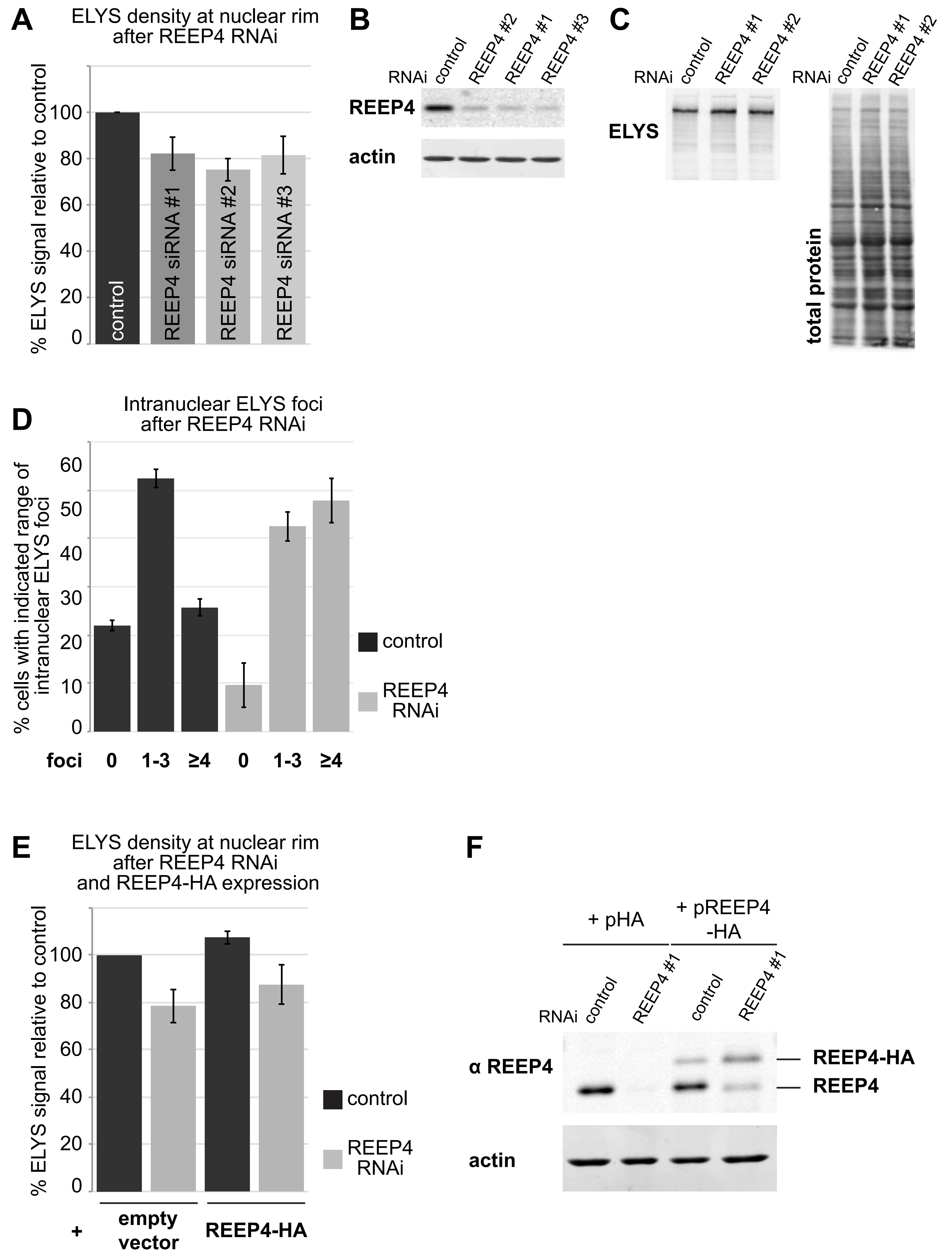
A
ELYS density at nuclear rim after REEP4 RNAi
% ELYS signal relative to control
0
20
40
60
80
100
control
REEP4 siRNA #1
REEP4 siRNA #2
REEP4 siRNA #3
B
RNAi
control
REEP4 #2
REEP4 #1
REEP4 #3
REEP4
actin
C
RNAi
control
REEP4 #1
REEP4 #2
ELYS
total protein
D
Intranuclear ELYS foci after REEP4 RNAi
% cells with indicated range of intranuclear ELYS foci
0
10
20
30
40
50
60
control
REEP4 RNAi
foci
0
1-3
≥4
0
1-3
≥4
E
ELYS density at nuclear rim after REEP4 RNAi and REEP4-HA expression
% ELYS signal relative to control
0
20
40
60
80
100
control
REEP4 RNAi
+
empty vector
REEP4-HA
F
+ pHA
+ pREEP4 -HA
RNAi
control
REEP4 #1
control
REEP4 #1
α REEP4
REEP4-HA
REEP4
actin

**Figure S1 (related to Figure 2). Characterization of BioID constructs.** HEK cells harboring the respective HA-TurboID-tagged constructs under an inducible promoter were induced for expression for 24 hours and treated with biotin for one hour before fixation and immunolabeling. TurboID constructs were detected with an anti-HA antibody, biotinylated proteins were stained with fluorescently labeled Streptavidin. Scale bar is 10 μm.

**Figure S2 (related to Figure 3). Role of ELYS in REEP4 INM targeting.** (A) Cells transfected with a construct for expression of REEP4-V5-tev-HA and with control, ELYS-, Nup153- or LBR-targeting siRNA were lysed in Laemmli buffer and analyzed by SDS-PAGE and immunoblotting using an antibody against REEP4, which detects both, endogenous REEP4 and REEP4-V5-tev-HA. Expression of REEP4-V5-tev-HA was not reduced after either of the siRNA treatments. Image on the right shows total protein stain to visualize the amounts of loaded protein in the different samples. (B, C, D) Lysates of cells transfected for the TEV assays with non-targeting control and (B) ELYS-, (C) Nup153- or (D) LBR-targeting siRNA were analyzed by SDS-PAGE and immunoblotting with antibodies against the respective depleted proteins. (B, C) ELYS and Nup153 protein levels were on average reduced by 70%. (D) LBR protein levels were on average reduced by 90%. (E) HeLa cells were treated with control or ELYS siRNA and immunostained for ELYS and the INM proteins Lamin A/C, SUN2, Emerin, and LBR. After ELYS-specific RNAi, ELYS was efficiently reduced (far left column) but localization of the INM proteins Lamin A/C, SUN2, and Emerin was not impaired. Lamin A/C staining appeared increased after ELYS depletion. Note that the two left columns show the same cells for control and ELYS RNAi, respectively, that were labeled for both, ELYS and LaminA/C. LBR targeting to the INM was impaired after ELYS depletion (far right column). Scale bar is 20 µm. (F) Cells expressing the INM protein Lap2β tagged N-terminally with HA-tev-V5 were treated with control or ELYS-targeting siRNAs and subjected to NusA-TEV treatment. Only the NusA-TEV treatment condition is shown, buffer-treated cells show an identical pattern. At least 30 cells were analyzed per condition in two different experiments. All control cells with V5-labeling, indicating successful transfection, also showed HA-staining of corresponding intensity. Among 68 V5-positive ELYS RNAi cells, one cell was lacking clear HA-staining but all other cells showed HA-staining that corresponded to V5 intensity, suggesting that ELYS depletion did not lead to increased permeability of NPCs for NusA-TEV protease. Scale bar is 10 µm.

**Figure S3 (related to Figure 4). Further characterization of REEP4 depletion phenotype.** (A) HeLa cells were transfected with control siRNA or one of three different REEP4 siRNAs (REEP4 siRNA #1 was used for the experiments shown in figure 4), fixed, immunolabeled for ELYS and imaged by confocal microscopy. Mean intensities at the nuclear rim were measured in control and REEP4 RNAi cells from the microscopy images. At least 100 cells were analyzed per condition, shown is the mean of three experiments. (B), (C) Whole lysates of HeLa cells depleted of REEP4 using the indicated siRNAs were analyzed by SDS-PAGE and Western blotting and probed for (B) REEP4 or (C) ELYS. To indicate amounts of loaded proteins, either (B) actin or (C) total protein were visualized. (D) The percentage of cells containing either “zero”, “one to three”, or “four or more” nucleoplasmic ELYS foci was determined in a blinded manner in control and REEP4 RNAi cells. Average of three experiments, at least 55 cells were analyzed per condition. (E, F) Empty HA-tagging vector or RNAi-resistant, HA-tagged wild-type REEP4 were expressed in control cells or REEP4 RNAi cells, fixed, immunolabeled for ELYS and imaged by confocal microscopy. ELYS mean intensity at the nuclear rim was measured in transfected cells identified by expression of co-transfected RFP-KDEL. Results of eight experiments are shown, at least 100 cells were analyzed per condition. Error bars are SEM. (F) Western blot analysis of lysates from cells used for an experiment described in (E). Endogenous REEP4 and REEP4-HA were detected with anti-REEP4 antibody, actin served as a loading control.
